# miR-146a and miR-200b alter cognition by targeting NMDA receptor subunits

**DOI:** 10.1101/2022.07.11.499553

**Authors:** Sowmya Gunasekaran, Ramakrishnapillai Vyomakesannair Omkumar

## Abstract

MicroRNAs (miRNAs) play pivotal roles in fine-tuning gene regulation. Understanding the mechanism of action of such miRNAs might help in manipulating the respective pathways thus providing therapeutic options. We have investigated the physiological roles of two miRNAs, miR-146a and miR-200b, that are differentially expressed in neurological disorders such as Alzheimer’s disease andschizophrenia.We specifically studied their involvement in learning and memory mechanisms. We show through bioinformatics prediction tools that these miRNAs can interact with transcripts of the N-methyl-D-aspartate receptor (NMDAR) subunits *Grin2A* and *Grin2B*. This was further supported by showing interaction of the miRNAs to the 3’UTR sequences of *Grin2A* and *Grin2B* through luciferase assay. Overexpression of these miRNAs in primary hippocampal neurons caused downregulation of GluN2B and GluN2A protein levels. Stereotactic injections of these miRNAs into rat hippocampus caused cognitive deficits in multiple behavioural tests along with decreased protein levels of the NMDAR subunits, GluN1, GluN2A and GluN2B, AMPAR subunit GluR1 and Neuregulin 1 (NRG1). During downregulation of NMDAR subunits by other physiological stimuli as in pharmacologically treated rat models [MK-801 treated and methylazoxymethanol acetate (MAM) treated], we found upregulated levels of miR-146a-5p and miR-200b-3p implying their involvement in downregulating NMDAR subunits. These results suggest the importance of miR-146a-5p and miR-200b-3p in mediating gene regulation in the hippocampus and their involvement in hippocampus dependent learning and memory.

## 1. INTRODUCTION

MicroRNAs (miRNAs) are small non-coding RNAs that regulate nearly 60% of the mammalian genes by post-transcriptional regulation (Friedman et al., 2009). They influence the stability and/or translation of the target mRNAs by binding to the 3’UTR sequence of the target mRNAs (Bartel, 2004; Ha and Kim, 2014). Thus, miRNAs fine tune the expression of various genes and orchestrate different functions of the nervous system (Rajman and Schratt, 2017). They play crucial roles in synaptic plasticity and memory in both vertebrates and invertebrates (Schratt, 2009). Differential expression of many miRNAs has been associated with neuropsychiatric and neurodegenerative disorders (O’Connor et al., 2016; MaciottaRolandin et al., 2013).Among the many miRNAs present in the brain, the mechanism of action is known for only a few (Sempere et al., 2004).

Glutamatergic signaling through N-methyl-D-aspartate receptor (NMDAR) is essential for several brain functions. NMDARs are excitatory ligand-gated ion channels, which are multiprotein complexes with two obligate GluN1 subunits and two GluN2 (A-D) / one GluN2 and one GluN3 (A and B) subunits. They play a prominent role in synaptic plasticity and synaptic pruning (Monyer et al., 1992).Their regulation and function are impaired in neuropsychiatric diseases such as schizophrenia and Alzheimer’s disease (Javitt, 2004).

There have been reports on miRNAs associated with the NMDAR pathway that act either on NMDAR or on targets upstream or downstream(Kocerha et al., 2009; Miller et al., 2012). miR-223 was shown to directly act on*Grin2B* using ischemic reperfusion brain injury models *in vitro* and *in vivo*. Genetic ablation of miR-223 leads to enhanced expression of GluN2B indicating it to be a potential molecule that protects against neuronal cell death in stroke and other excitotoxic conditions (Harraz et al., 2012). miR-137, a candidate gene in schizophrenia, regulates GluN2A and thus alters synaptic plasticity and imposes an inhibitory effect on post stroke depression (Zhao et al., 2013). miR-19a and miR-539 can also target and regulate NMDAR (Corbel et al., 2015).In our previous study, we found miR-129-2, miR-148a and miR-296 to be targeting NMDAR*in vitro*and inanimal models in which NMDAR is dysregulated (Gunasekaran et al., 2021).

In this study we have investigated whether miR-146a and miR-200b, which are differentially expressed in diseases such as Alzheimer’s disease and schizophrenia (Beveridge et al., 2010; Perkins et al., 2007; Hall et al., 2015; Ghazaryan et al., 2019; Ibrahim et al., 2020;Moreau et al., 2011;Santarelli et al., 2011; Wang et al., 2016; Mai et al., 2019; Liu et al., 2014;Fu et al., 2019), can regulate NMDAR subunits. We established the interaction of these miRNAs with NMDAR subunits through bioinformatic analysis and luciferase assay. We obtained further validation on these miRNAs by showing their ability to downregulate NMDAR subunits upon overexpressing them in primary hippocampal neurons. We also injected these miRNAs to hippocampus *in vivo* to understand the physiological role of these miRNAs through behavioural and subsequent biochemical experiments. In addition, we also investigated the levels of these miRNAs in pharmacological rat models in which NMDAR is known to be downregulated.

## 2. MATERIALS AND METHODS

### 2.1 Materials

β-mercaptoethanol, 30% acrylamide,ammoniumpersulphate (APS), dithiothreitol (DTT), Tween-20,Tris,protease inhibitor cocktail (PIC),sodium dodecyl sulphate (SDS),sodium hydroxide (NaOH), phenylmethylsulfonyl fluoride (PMSF), MK-801 (dizocilpine), tetramethylethylenediamine (TEMED), Calcium chloride (CaCl2), Taq DNA polymerase and phusion DNA polymerasewere purchased from Sigma (USA). DNA ladders, T4 DNA ligase, and enzymes for restriction digestion were purchased from New England biolabs (USA) orFermentas (USA). Clarity ECL reagent, nitrocellulose membrane and prestained protein marker were purchased from Bio-Rad Laboratories (USA). Ampicillin, LB agar,bovine serum albumin (BSA), agarose and LB broth were purchased from GE Healthcare Life Sciences (USA). Methylazoxymethanol acetate (MAM) was purchased from FUJIFILM Wako Pure Chemical Corporation (Japan). Dulbecco’s modified eagle medium (DMEM), neurobasal medium (NBM), fetal bovine serum (FBS), glutamax, Pierce BCA protein assay kit, antibiotic-antimycotic, trypsin, mirVana kit, lipofectamineand lipofectamine2000 reagent, Opti-MEM medium, Hank’s balanced salt solution (HBSS), B-27 supplement and DNAse I were purchased fromThermo Fisher Scientific (USA). High-Capacity cDNA Reverse Transcription Kit and Mir-X™ miRNA First Strand Synthesis Kit were purchased from Applied Biosystems (USA) and Takara (China) respectively. Antibodies used for western blotting were purchased either fromSanta Cruz Biotechnology (USA), Sigma (USA),Abcam (UK), Millipore (USA) or Cell Signaling Technologies (CST) (USA) for GluR1 (Cat. No-Sc55509), GluN2B (Cat. No-Ab65783 or sc9057),NRG1 (Cat. No-Sc28916), GluN2A (Cat. No- #4205S or MAB5216),β-actin (Cat.No-A53l6)and GluN1 (Cat. No-ab17345). Adeno-associated virus (AAV) was purchased from Applied Biological Materials (abm). AAV-mir-GFP-Blank Control Virus (Serotype 8) (Cat. No-Am00108) (Abbreviated as VC for vector control), GFP rno-mir-200b AAV miRNA Virus (Serotype 8) (Cat. No-Amr1010308)(Abbreviated as AAV-miR-200b) and GFP rno-mir-146a AAV miRNA (Cat. No-Amr1005808) (Abbreviated as AAV-miR-146a) were procured.

### 2.2 Methods

#### 2.2.1 Bioinformatic analysis

The sequences of the miRNAs were obtained using miRBase database (http://www.mirbase.org/) and the sequences of the 3’UTR of *Grin2A/Grin2B* was obtained form UCSC Genome browser (https://genome.ucsc.edu/).

The tools used for mining the miRNAs that target the genes *Grin2A* and *Grin2B* wereTargetScan (http://www.targetscan.org/vert_72/), miRanda (http://www.microrna.org/), MicroCosm (https://tools4mirs.org/software/mirna_databases/microcosm-targets/), Segal/PITA (https://genie.weizmann.ac.il/pubs/mir07/mir07_prediction.html), miRDB (http://mirdb.org/) and RNAhybrid tool (https://bibiserv.cebitec.uni-bielefeld.de/rnahybrid/). We chose only those miRNAs, which were predicted in more than three tools to negate false positives.

#### 2.2.2 DNA constructs

UCSC database (https://genome.ucsc.edu/) was used for obtaining precursor miRNA (pre-miRNA) sequences for miR-146a and miR-200b. Suitable primers were designed (Table 1) and were used for cloning these pre-miRNAs from rat genomic DNA. The amplified pre-miRNAs were cloned into the expression vector, pRIPM that has Ds Red as marker.. The 3’UTRs of *Grin2A* and *Grin2B* in the vector backbone psiCHECK2 described before (Gunasekaran et al., 2021) were also used for performing the dual luciferase assay. All the clones were confirmed by restriction digestion and sequencing.

**Table 1.**
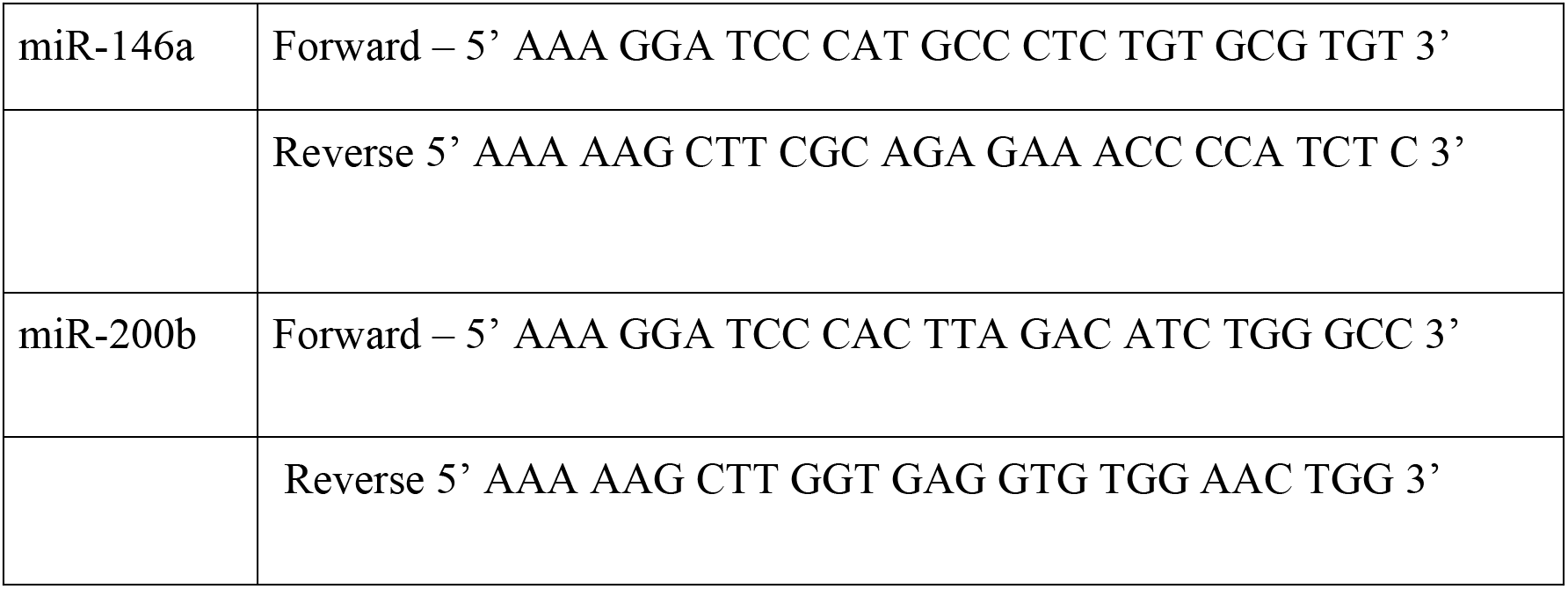
Primers used for cloning pre-miRNAs. Primers used for cloning the pre-miRNAs into the pRIPM vector were designed using the UCSC genome database.

#### 2.2.3 Luciferase assay

The procedure for dual luciferase assay was performed using Human Embryonic kidney-293 (HEK-293) cells as described before (Gunasekaran et al., 2021). In brief, The psiCHECK2 and pRIPM vector constructs carrying the 3’UTRs and the pre-miRNAs respectively were used for the assay. These vectors were cotransfected and after 48hrs reported assay for performed. The empty pRIPM vector along with *Grin2A/Grin2B* 3’UTR psiCHECK2 were used as control for this experiment.

#### 2.2.4 Animals

We used Wistar rat strain for our study. For primary cultures of hippocampal neurons, rat embryos of embryonic day 18-19 (E18-19) were used. AAVs carrying the pre-miRNAs were streotactically injected into Male Wistar rats of adult stage (60-70 day old, 200-300g). For the MK-801 model, and male rats of adolescent stage (40-45 day old, 100-150g) were used. For generating the MAM model, female dams were used. All the animals were housed with 12 hr light-dark cycle. Access to chow and water were ad libitum to the animals. The procedures followed the rules and regulations of the Committee for the Purpose of Control and Supervision of Experiments on Animals (CPCSEA), Government of India. The protocol was approved by the Institutional Animal Ethics Committee (IAEC) of Rajiv Gandhi Centre for Biotechnology.

##### 2.2.4.1 Primary neuronal culture and Transfection

Pregnant female dams of E-18-19 were sacrificed and the hippocampus was dissected, trypsinised and the dissociated cells were cultured in coated dishes. The primary neurons were maintained up to 20 days *in vitro* (DIV 20) (Gunasekaran et al., 2021;Chandran et al., 2021; Salazar et al., 2017).

Neurons were transduced using AAVs containing the pre-miRNAs, miR-146a (AAV-miR-146a), miR-200b (AAV-miR-200b) or vector control (VC) on DIV 7. The viral titre for all the AAVs were 10^8^ GC/ml. The vector map for these constructs are shown in Suppl. Fig 2. The cells were harvested on DIV 18-20 and were further processed for conducting western blotting experiments.

##### 2.2.4.2 Treatment regimes for the animal models

###### 2.2.4.2.1 Stereotactic injections of AAV to male Wistar rats

The rats were deeply anesthetized by isoflurane (2-4%) inhalation. The head was mounted on a stereotaxic frame (Company name: BenchMarkTM, Coretech Holding Scientific, USA currently merged to Leica Microsystems, Germany). The coordinates used for bilateral intrahippocampal injections at the CA1 site were: AP=-3.8 mm, ML=+1.4 mm and DV=-2.7 mm (Paxinos et al., 1980).All AAV constructs carried EGFP as marker. The viral titre used was 1X 10^8^ GC/ml and around 5 μL was injected per side using a 25 μL Hamilton syringe. The solution was injected very slowly and the needle was kept undisturbed for another 2 min to avoid backflow of the fluid. Sham control was used with PBS injection. The animals were kept for 4-5 weeks for recovery followed by behavioural tests. These animals were then sacrificed by cervical dislocation and the hippocampal region was dissected to measure the protein and mRNA levels of different targets as well as miRNA levels using western blotting and qPCR.

###### 2.2.4.2.2 MK-801 model and MAM model

MK-801 (0.5 mg/Kg) was dissolved in saline and was injected intraperitoneally for 5 consecutive days (one injection a day) at consistent time period of 2 pm-5 pm to male adolescent rats. MAM (20 mg/Kg) was dissolved in saline and was injected by intraperitoneal route to pregnant dams on gestational day 17 (GD 17). Weaning of the pups was performed on postnatal day 30 (P30). Detailed procedure was described before (Gunasekaran et al., 2021).

The experimental rats were subjected to behavioural testing to confirm impairments, sacrificed by cervical dislocation and the hippocampi were dissected for performing biochemical analysis.

##### 2.2.4.3 Behavioural analysis

Open field test (OFT), novel object recognition (NORT), object location tests (OLT) and Morris water maze (MWM) test were used for behavioral analysis as described before (Gunasekaran et al., 2021).

The animals were made to acclimatize to the behavior chamber one hour prior to the start of each experiment.

In brief, OFT was carried out in an open square box. The total distance traveled and the time spent in central/peripheral zone by the animal in the box for 10-11 min was recorded.

NORT was also conducted in the open square box with 2-3 non-toxic objects. The initial habituations for the animals were carried out for 10 min in the empty arena. After habituation period of 24hrs hours, the animal was given training in the familiar arena where they were exposed to two identical non-toxic objects for 5 min. During the test period which is 1 hr and 24 hrs after the training period, the animals were exposed to one novel object and one familiar object for 5min in the same arena. The time taken for exploring the novel and familiar object were noted and accordingly discrimination index (DI) and recognition index (RI) were calculated.

In OLT, the animal was subjected to training wherein it was made to explore two familiar objects for 5 min in the open square box. After one hour, the animal was put in the same arena with one of the objects displaced to a new location. This is the test period, which was done for 5 min and the time taken to explore the displaced object and the familiar object by the animal were noted. DI and RI were calculated.

The water maze test was performed in a white tank having a small circular platform inside. The whole tank was filled with water which was translucent enough to make the platform was invisible. The animals were trained for 5 days to locate the hidden platform using learning cues. Each day, the animals underwent five trials of 1 min each with intertrial intervals of 20-30 seconds. The time taken to reach the platform was noted and the average escape latency was calculated for each animal on each day.

NoldusEthoVision XT software (Noldus Information Technology, Wageningen, The Netherlands) was used for analyses of the video files. After the MWM test, animals were sacrificed and the brains were immediately removed, dissected, snap frozen and stored in −80°C.

##### 2.2.4.4 Western Blotting

The hippocampal neurons were scrapped and were washed with 1X PBS. The cell pellet was lysed using RIPA lysis buffer with 50 mM β-glycerophosphate, 50 mM sodium fluoride, 200 μM sodium orthovanadate and 5 mM ethylenediaminetetraacetic acid (EDTA). The hippocampal tissues from animals were homogenized using liquid nitrogen followed by solubilizing them in lysis buffer [150 mM NaCl, 1 mM dithiothreitol (DTT), 1% NP-40, 50 mM Tris-HCl, pH 7.4, 5 mM EDTA, 10 mM staurosporine, 50 mM β-glycerophosphate, 5 mM sodium orthovanadate, 1% SDS, 0.2 mM phenylmethylsulphonylfluoride (PMSF), and 1X complete protease inhibitor cocktail].Bicinchoninic acid (BCA) method was used to quantitate the protein concentration in the sample and 60-100 μg of protein were subjected to SDS-PAGE for each sample. The whole procedure was carried out as described before (Gunasekaran et al., 2021). After transfer procedure, the membrane has been cut horizontally. Different molecular size regions were processed separately. The blots were incubated overnight first with the corresponding primary antibodies. The primary antibodies used were: rabbit anti-GluN1 (1:1000, Abcam), rabbit anti-GluN2B (1:1000, Abcam), mouse anti-GluN2A (1:1000, Millipore), mouse anti-GluR1 (1:750, Santacruz) mouse anti-beta actin (1:3000Sigma) and rabbit anti-NRG1 (1:1000, Santacruz). Subsequently, the blots were incubated for 2 hrs at room temperature with secondary antibodies, which are horseradish peroxidase (HRP)- conjugated (1:3000 or 1:10,000-Sigma). The blots were developed using Clarity ECL reagent in Chemidoc gel apparatus (Biorad, USA). The band intensity was quantitated using ImageJ software and the band intensity of β-actin was used for normalization.

##### 2.2.4.5 Real time PCR analysis of hippocampal tissues

Extraction and measurement of transcripts (mRNAs and miRNAs) in the brain tissues were carried out by Quantitative real-time PCR (qPCR) as described before (Gunasekaran et al., 2021) according to manufacturer’s instructions.

KiCqStart SYBR green primers (Sigma) were used for quantitating the mRNA levels (Table-2 shows the primers used for quantitating miRNAs and mRNAs in. From the 2^ΔCT values, relative expression levels were calculated in each sample. Data presented using the average value from minimum three independent experiments and β-actin was used for normalisation.

**Table 2.**
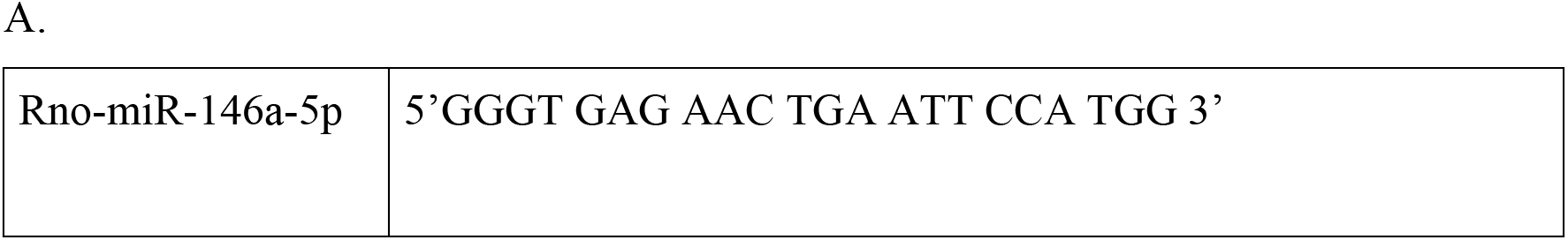
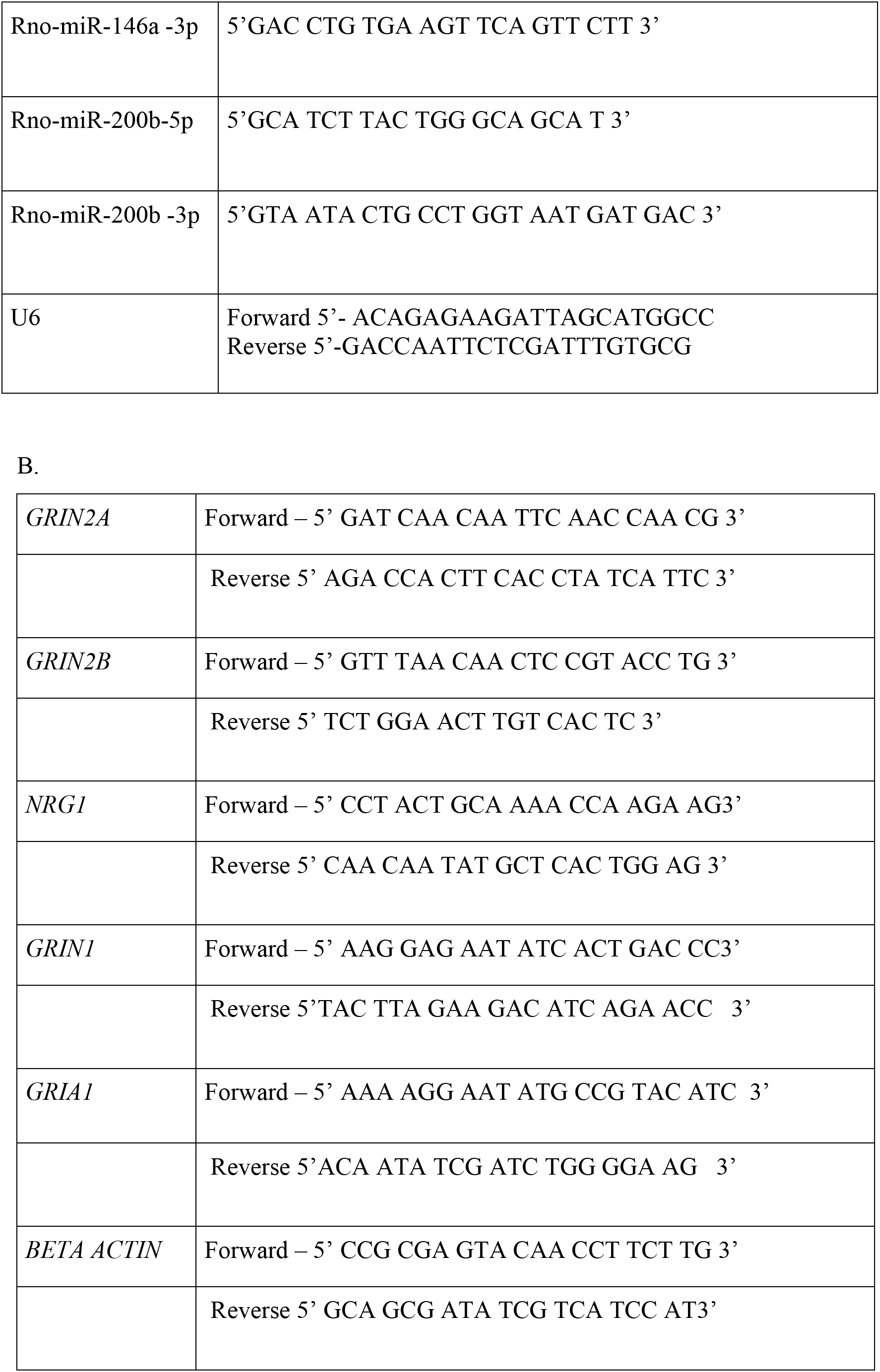
Primers used for real time PCR of miRNAs and genes. A: Primers used for amplifying the miRNAs. B: Primers used for amplifying the transcripts of the genes

#### 2.2.5 Statistical Analysis

The data is presented asmean ± standard deviation (SD) for independent experiments/replicates. Luciferase assay data (n=4) was analyzed using parametric unpaired Student’s t test between each miRNA group and the corresponding control group. Behavioral data analysis (n=6-8 per group) for stereotactic injection experiments was done using one-way ANOVA followed by Dunnett’s post hoc test between the control and the treatment groups. Data from the MWM tests were analyzed using repeated measures two-way ANOVA followed by Dunnett’s post hoc test. The data from the biochemical analysis for AAV experiments (n=4-5) for qPCR and western blotting was done using one-way ANOVA followed by Dunnett’s post hoc test. The statistical analysis for the behavior experiments was completely blinded. Data from the qPCR (n=3-5 per group) for MK-801 and MAM model were statistically analyzed using unpaired parametric Student’s t test. The data was analyzed using the GraphPad prism 8 software (8.4.2). Statistical significance was set at p<0.05.

## 3. RESULTS

### 3.1. miR-146a and miR-200b bind to 3’UTRs of NMDAR subunits

To find the miRNAs that bind to the 3’UTRs of the NMDAR subunits *Grin2A* and *Grin2B*, we used 3-5 online prediction tools and found numerous hits. We chose miR-146a and miR-200b for further studies because they were found to be differentially expressed in neurodegenerative disorders such as Alzheimer’s diseaseand also in neuropsychiatric disorders such as schizophrenia (Wang et al., 2016; Mai et al., 2019; Liu et al., 2014;Fu et al., 2019; Beveridge et al., 2010; Perkins et al., 2007; Hall et al., 2015; Ghazaryan et al., 2019; Ibrahim et al., 2020;Moreau et al., 2011;Santarelli et al., 2011). miR-146a-5p has binding sites on *Grin2B* and miR-200b-3p has binding sites on *Grin2A* with rat query sequence. Among the binding sites, at least one has strong conservation among other vertebrates including the humans (Figure 1B and 1C). The converse pairing of miR-146a-5p with *Grin2A* and miR-200b-3p with *Grin2B* are also possible at different binding sites but the sites are conserved only in mouse, rat and squirrel or mouse and rat respectively (Figure 1A and 1D). The site type was 7mer-m8 or 7mer-A1 or 8mer, implying strong binding of the miRNA seed sequence to the 3’UTR of the target. In case of the complimentary strands of the miRNAs, miR-146a-3p has binding sites on *Grin2A* and *Grin2B* and miR-200b-5p has sites on *Grin2A*, but the sites are not conserved among other species including humans (data not shown). We used RNA hybrid tool, that uses minimum free energy (mfe) paradigm to understand the binding stability of the miRNA-mRNA interaction (Figure 1 A-D).

**Fig 1.**
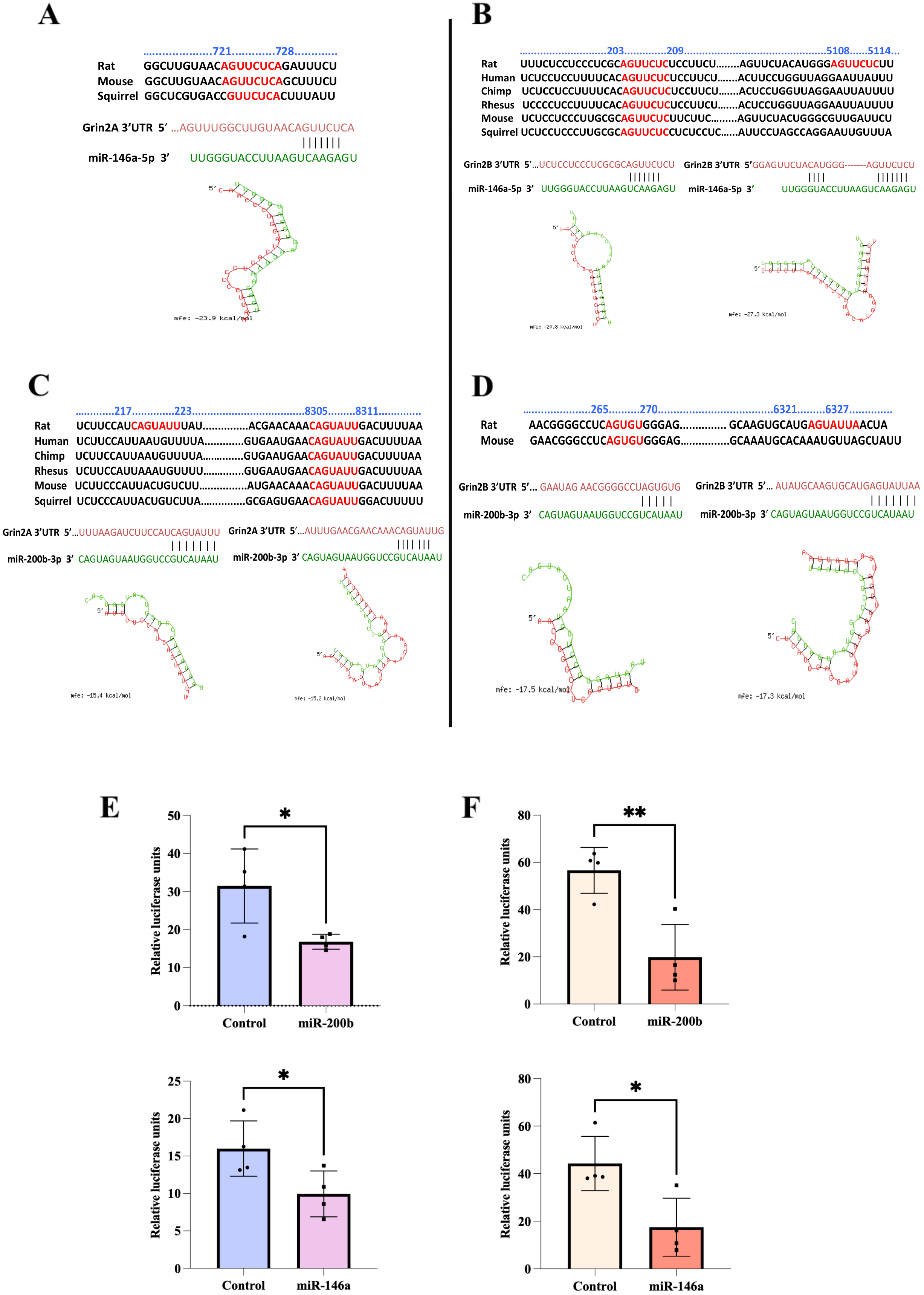
Interactions of miR-146a-5p and miR-200b-3p with *Grin2A/Grin2B* 3’UTR. Sequence alignment of the binding sites of miR-146a-5p (A and B) and miR-200b-3p (C and D) on*Grin2A* (A and C) and *Grin2B* (B and D) indicating conservation among different species is shown. Graphics illustrates the predicted binding of miRNA sequence (green) with the target 3’ UTR sequence (red) obtained using RNA hybrid tool with corresponding minimum free energy (mfe) shown (A-D). E and F show the dual luciferase assay results showing interaction of the miRNAs with the 3’UTR sequences *in vitro*. HEK-293 cells were cotransfected with psiCHECK2 plasmids carrying the 3’UTR of *Grin2A*/*Grin2B* along with pRIPM Ds Red plasmids carrying the pre-miRNA sequences of miR-146a and miR-200b. Control represents co-transfection of the psiCHECK2 plasmids having 3’UTRs with empty pRIPM Ds Red plasmids without the pre-miRNA sequence. The firefly luciferase activity arising from the psiCHECK2 vector was used for normalization and the renilla luciferase activity was used as the reporter of interaction. Statistical analysis was done by unpaired Student’s t test, **p ≤0.01, *p ≤0.05, n=4. Data are expressed as mean ±SD.

To experimentally validate the predicted miRNA-mRNA interactions, we performed dual luciferase assay. We constructed plamsids with 3’UTR sequences of *Grin2A* and *Grin2B (Rattus norvegicus*) in psiCHECK2 vector. The pre-miRNA sequences for miR-146a and miR-200b were cloned in pRIPM DsRed vector. Each pre-miRNA was cotransfected with *Grin2A* or *Grin2B* 3’UTR in HEK-293 cells and after 48 hrs the cells were subjected to the assay. The control group had co-transfection of empty pRIPM DsRed vector with *Grin2A* or *Grin2B* 3’UTR vector. It was observed that both miR-146a and miR-200b showed significant reduction in the renilla luciferase activity against *Grin2A* 3’UTRimplying strong interaction [miR-200b (t =2.95, df=6); miR-146 (t=2.52, df=6)] (Figure 1E). Similarly both the miRNAs showed significant interaction with *Grin2B* 3’UTR [miR-200b (t =4.34,df=6); miR-146 (t=3.20, df=6)] (Figure 1F). Theseresultsshow that the predicted miRNAs can interact with functional binding sites in the 3’UTRs of *Grin2A* and *Grin2B*.

### 3.2. Downregulation of the NMDAR subunits by overexpression of miR-146a or miR-200b in primary hippocampal neurons

We further wanted to investigate whether the miRNA-mRNA binding described above could cause alteration in GluN2A/GluN2B protein levels in a neuronal system. For this we used primary hippocampal neurons prepared from E-18 rat embryos. The cultures were subjected to overexpression of the miRNAs by treatment with the corresponding AAV constructs, AAV-miR-146a,AAV-miR-200b orVC on DIV 7 and the cells were harvested on DIV 18-20. The transduction efficiency was more than 70% (Figure 2A) as seen from the expression of the EGFP marker encoded in the viral constructs (Suppl. Figure 2). We checked for protein expression by western blotting and found significant reduction of GluN2A and GluN2B in neurons transduced by either miR-146a or miR-200b (Figure 2B and C). Analysis using one-way ANOVA showed significant changes among the three groups using the Dunnetts test [F (2,6)=16.66, P=0.0036 for GluN2A; F (2,6)=14.61, P=0.005 for GluN2B) (Figure 2B and C). We also found decrease in GluN1 subunit of the NMDAR implying that the miRNAs caused downregulation of all the major subunits of NMDAR (Figure 2D). The one-way ANOVA showed significant difference between the groups for GluN1 [F (2,6)=5.13, P=0.050]. We checked for the other targets of these miRNAs such as neuregulin1 (NRG1) and found significant downregulation by miR-200b [F (2,6)=10.88, P=0.01] (Figure 2E). Overall, these results convey that miR-146a and miR-200b can downregulate NMDAR subunits GluN2A and GluN2B *in vitro* along with downregulating other proteins such as GluN1 and NRG1.

**Fig 2.**
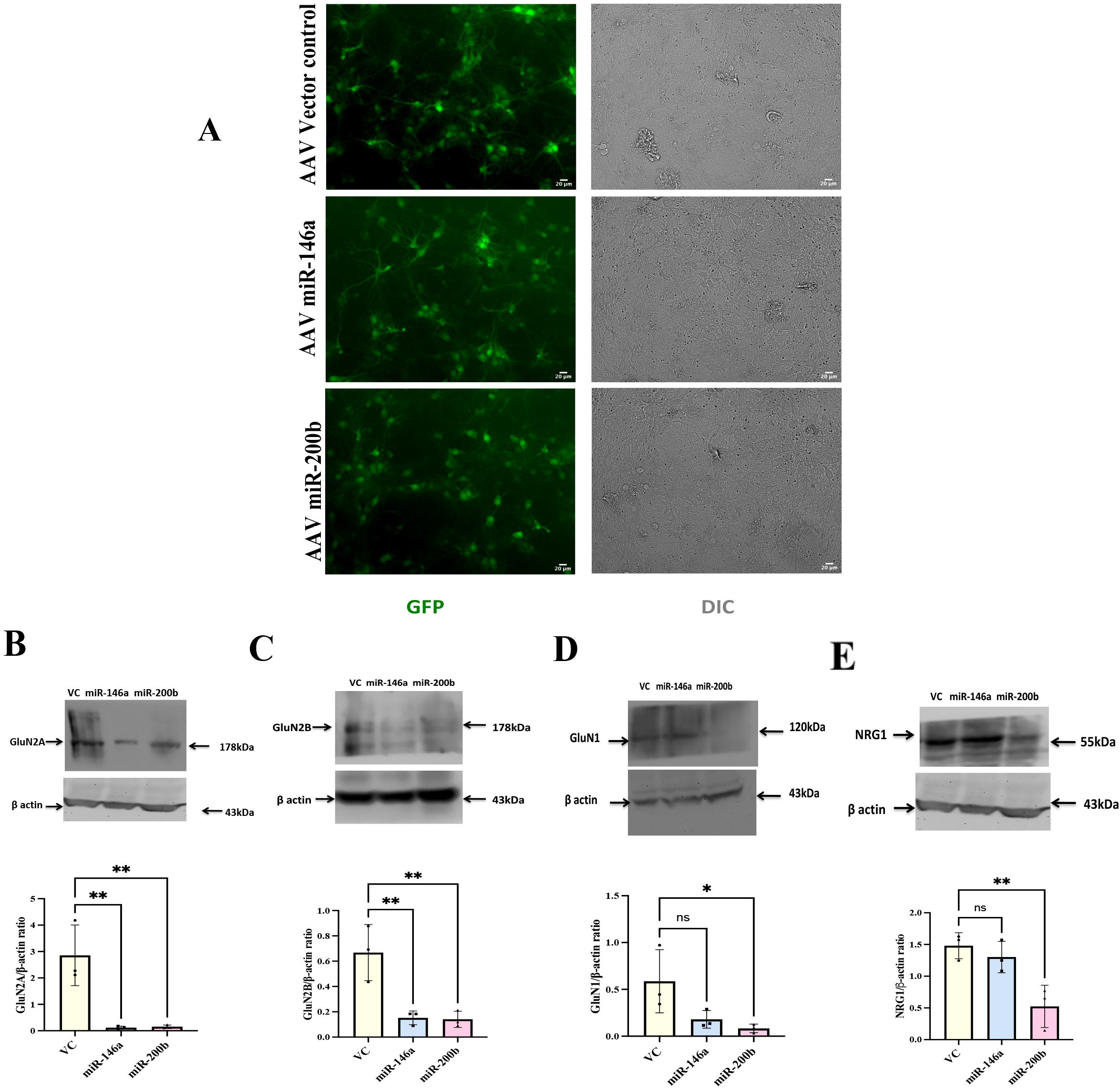
miR-146a and miR-200b downregulate GluN2B and GluN2A levels in primary hippocampal neurons. The hippocampal neurons were transduced with AAV-miR-146a and AAV-miR-200b and VC on DIV 7, harvested on DIV 18-20 and were processed for western blotting. A: Representative images showing the transduction efficiency of the neurons on DIV 20. B-E: Representative western blots for GluN2A, GluN2B, GluN1and NRG1. Since GluN2A and NRG1 were probed in the same blot at different molecular size regions, they share the same actin blot. Quantitation is done using Image J software. Data are expressed as mean ±SD. The target protein band intensity is normalized toactin band intensity for multiple samples and is shown along with the respective immunoblots. Statistical analysis was done using one-way ANOVA with Dunnett’s test. **p ≤ 0.01, *p ≤ 0.05; n=3.

To support the western blot data, we also studied expression of GluN2B in primary neurons by immunocytochemical staining after overexpressing the miRNAs. The pRIPM-Ds Red vector construct carrying pre-miR-146a or pre-miR-200b was transfected to cultures at DIV 7. The cells were fixed on DIV 10 and were subjected to immunocytochemical staining. We found decrease in GluN2B expression in miR-146a and miR-200b transfected cells (Suppl Figure 1) [miR-146a (t=3.70, df=38), miR-200b (t=2.23, df=38)]. The Ds Red present in the constructs marked the transfected neurons. The empty pRIPM-Ds Red vector was used as control for this experiment, which did not show any detectable GluN2B downregulation (Gunasekaran et al., 2021).

### 3.3. Regulation of NMDAR by miR-146a and miR-200b *in vivo*

To evaluate the potential role of miR-146a and miR-200b *in vivo*, we performed stereotactic surgeries tointroduce these miRNAs into rat hippocampus. Adult male rats were bilaterally injected with AAV containing the pre-miRNAs (miR-146a or miR-200b) at the CA1 region of the hippocampi. Empty vector (VC) and sham injection (using 1X PBS) were used as controls. The animals were allowed to recover for 4-5 weeks followed by behavioural and biochemical experiments. The treatment regime used is shown in Figure 3A. Presence of the virus in the CA1 region was shown by expression of the EGFP marker protein, which is present along with the miRNA sequence in the vector backbone (Suppl. Figure 2). Figure 3B shows the representative image of AAV-mir-200b injected hippocampus, 4-5 weeks after injection.

**Fig 3.**
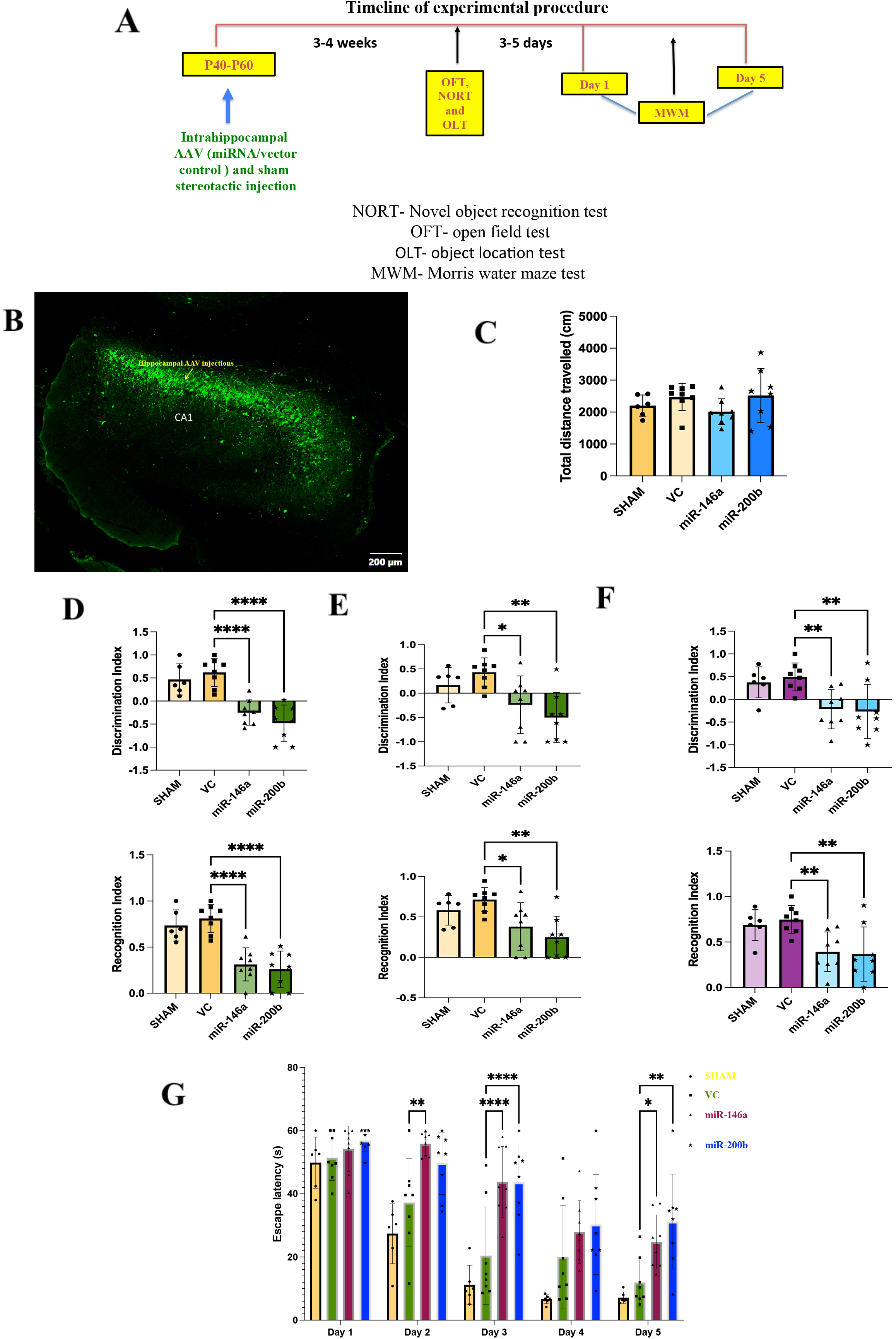
Animals injected with miRNAs show impaired learning and memory. A: Timeline of the experimental procedures. B: Representative fluorescence microscopy image of a coronal section of the brain of rat 4 weeks after stereotactic injection in the CA1 area of hippocampus with AAV-mir-200b particles. The EGFP present in the construct serves as a marker and allows to verify the site and the efficacy of the injection. The presented image was obtained by automatically aligning and stitching 50 tiled images into a mosaic (using Olympus confocal microscope). C-G: Behavioural analysis of the animals, four weeks after stereotactic injections of AAV-miR-146a orAAV-miR-200b or VC constructs. Sham injections were done using 1X PBS solution. C: OFT to measure the total distance travelled inside the box for 10-11 min. D and E: Results of NORT showing DI and RI, 1 hr (D) and 24 hrs (E) after training. F: Results of OLT conducted 1 hr after training. Quantitation is shown as mean ±SD. Statistical analysis was done by one-way ANOVA followed by Dunnett’s post hoc multiple comparison test with p values indicated: ****p ≤0.0001, **p ≤ 0.01, *p ≤ 0.05, n=6 per group for sham, n=8 per group for VC, AAV-miR-146a and AAV-miR-200b. G: MWM test showing the latency to reach platform for 5 days. Quantitation is shown as mean ±SD. Statistical analysis was done by repeated measures two-way ANOVA with Dunnett’s multiple comparison test with p values indicated: ****p ≤0.0001, **p ≤ 0.01, *p ≤ 0.05; n=6 per group for sham, n=8 per group for VC, AAV-miR-146a and AAV-miR-200b.

#### 3.3.1. miR-146a and miR-200b injected animals show cognitive disability

The animals were first subjected to OFT to assess the anxiety levels and locomotor ability. It was observed that the animals injected with AAV-miR-146a and AAV-miR-200b performed similar to sham and VC, implying no major impairment in locomotor activity (Figure 3C). The miRNA injected animals did not seem to have increased anxiety as observed by the percentage of time spent in the peripheral and central zone (PZ and CZ), which showed no significant differences among the groups (Suppl Fig 3).

The animals then underwent tests for learning and memory-NORT and OLT. It was observed that in the NORT tests done 1 hr and 24 hrs after training, both the miRNA injected groups had significant reduction in the discrimination index (DI) and recognition index (RI) when compared to VC (Figure 3D and 3E) indicating that the miRNA injected animals did not spend more time in exploring the novel object, implying memory deficits. The one-way ANOVA showed significant differences between the groups for tests done 1 hr [DI (F _(3,26)_ = 20.31; p<0.0001) and RI (F_(3,26)_ = 19.9; p<0.0001)] and 24 hrs after training [DI (F_(3,26)_ = 6.18; p=0.0026) and RI (F_(3,26)_ = 6.18; p=0.0026)].

In OLT, the miRNAinjected animals did not spend more time in exploring the displaced object when compared to VC. RI and DI were significantly reduced in the miRNA injected animals than VC indicating memory impairment (Figure 3F). The one-way ANOVA analysis showed significant difference among the groups [DI (F _(3,26)_ = 6.07; p=0.0028) and RI (F _(3,26)_ = 6.07; p=0.0028)].

Lastly, Morris water maze experiments were conducted to check the hippocampal dependent spatial learning and memory (Gathe and Kempermann, 2013). The miRNA treated animals showed significant learning impairment on day 3 and day 5 (Figure 3G). Although performance appears impaired on day 4, the data was not statistically significant. Animals injected with AAV-miR-146a showed significant learning impairment even on day 2 of the experiment. The repeated measures two-way ANOVA indicated significant differences between the time and groups (F _(12,130)_ = 1.97; p=0.031).

These results showed that the animals injected with AAV-miR-146a or AAV-miR-200b had severe cognitive impairment with learning and memory deficits. Because there was no significant difference between sham and VC animals in the behavior experiments, only VC animals were used as controls for subsequent miRNA, mRNA and protein analysis.

#### 3.3.2. Biochemical analysis indicates miR-146a and miR-200b downregulate NMDAR via translational repression in hippocampus

The 5p and 3p forms of both miR-146a and miR-200b were significantly high in the animals injected with the AAV constructs of the respective miRNAs (Figure 4A). The one-way ANOVA analysis showed significant differences among the groups injected with the AAV constructs [F (2,11) = 9.96; p=0.003for miR-146a-3p; F _(2,11)_ = 7.53; p=0.008for miR-146a-5p; F _(2,11)_ = 6.55; p=0.013for miR-200b-3p; F _(2,11)_ = 7.81; p=0.007for miR-200b-5p]. As expected, the level of only the injected miRNA increased and the other miRNA was not significantly altered in the respective experimental group (Figure 4A).

**Fig 4.**
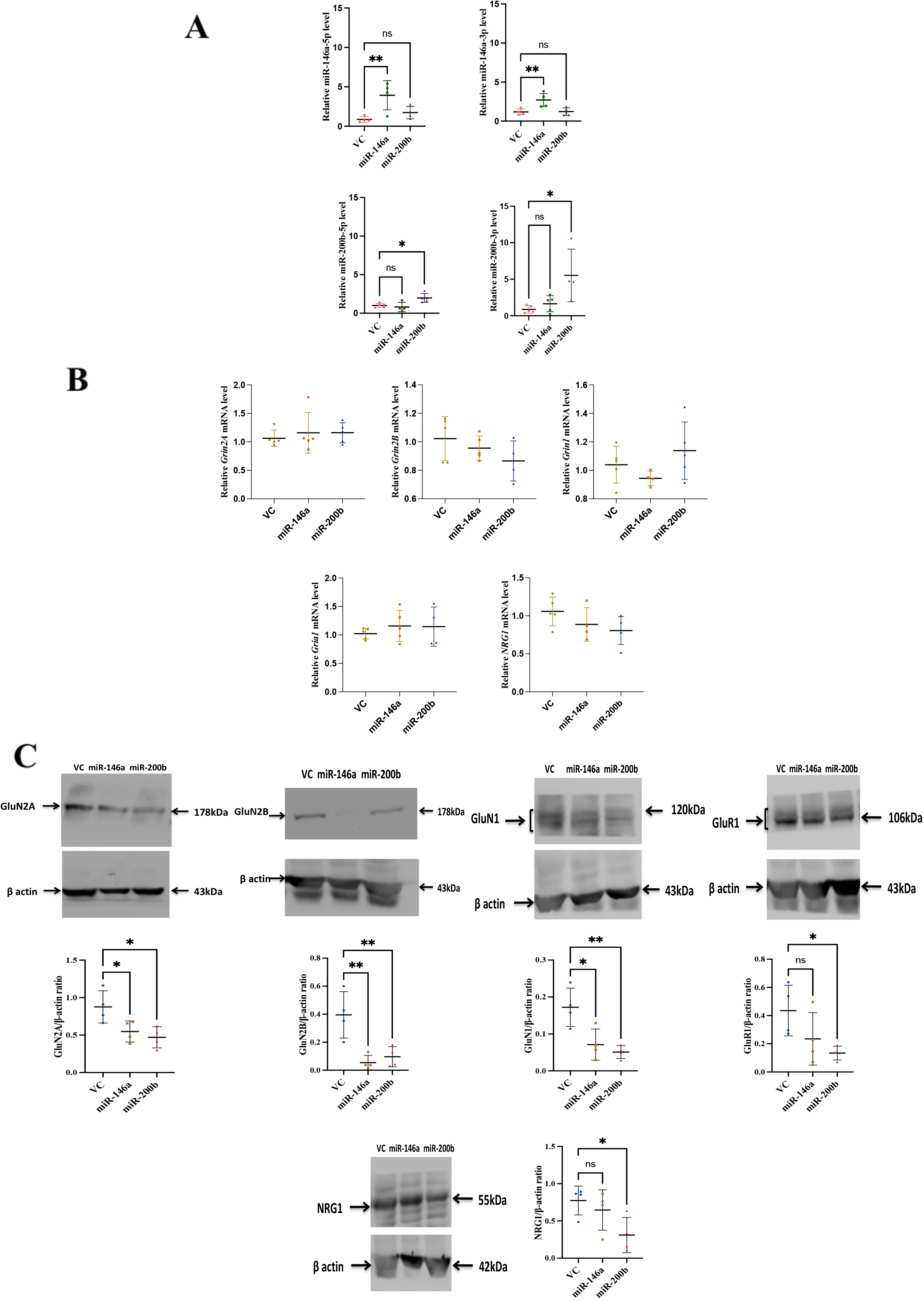
Alternations in miRNA, mRNA and protein levels in the hippocampal region of the animalsinjected with AAV particles. A: Expression levels of miR-146a-5p, miR-146a-3p, miR-200b-5p and miR-200b-3p in animals injected with AAV as estimated by real time PCR. B: Quantitation of transcript levels done by real time PCR for *Grin2A*, *Grin2B, Grin1, Gria1* and *NRG1*. C: Protein expression levels of GluN2A, GluN2B, GluN1, GluR1 and NRG1. Representative western blots along with quantitation is shown. Quantitation is done using ImageJ software. The target protein band intensity is normalized to actin band intensity for multiple samples and is shown along with the respective immunoblots. Statistical analysis was done by one-way ANOVA with Dunnett’s post hoc test. Data is represented as mean ±SD. **p ≤ 0.01, *p ≤ 0.05; n=4-5.

The transcript levels for the proteins that were downregulated by the miRNAs in hippocampal neurons in culture (Figure 2), were checked in the miRNA injected animals. The NMDAR subunit transcripts, *Grin2A* and *Grin2B*, that are targets of miR-146a and miR-200b, were not significantly altered when compared to the control groups (Figure 4B). The levels of *NRG1*also were largely unchanged, implying that the injected miRNAs are not altering the transcript levels of the target proteins. There was no significant change in the levels of *Grin1* transcripts (Figure 4B). We also checked the expression of AMPA receptor *Gria1*due to impaired memory in the miRNA overexpressed animals and found no change in the transcript level.

The protein levels of these transcripts were analysed and it was found that GluN2A and GluN2B levels for both the miRNA injected groups were significantly reduced (Figure 4C). The one-way ANOVA showed significant differences between the groups (F _(2,9)_ = 6.43; p=0.018 for GluN2A; F _(2,9)_ = 11.72; p=0.003 for GluN2B). NRG1 protein was found to be significantly downregulated in the miR-200b injected animals. GluN1 was also found to be significantly reduced in both the miRNA injected groups with one-way ANOVA showing significant difference between the groups (F (2,9) = 10.58; p=0.004). GluR1 was significantly decreased in AAV-miR-200b injected animals.

These results indicate that NMDAR subunits GluN2A and GluN2B are downregulated by miR-146a-5p and miR-200b-3p, most likely by translational repression without altering the levels of their transcripts.

#### 3.3.3 Altered expression levels of miR-146a and miR-200b in pharmacological models with NMDAR downregulation

The data so far shows that miR-146a and miR-200b can downregulate NMDAR subunits *in vitro* and *in vivo*, upon overexpression. We then checked if these miRNAs play a role in downregulating NMDARs in other physiological conditions of NMDAR hypofunction.For this we chose two pharmacological rat models in which NMDAR is downregulated by physiological signals that are indirectly induced by the experimental treatment (Coyle, 2012).We used the MK-801 model and MAM model created as reported before (Gunasekaran et al., 2021). MK-801 is an NMDAR antagonist, and is used for creating a pharmacological model by a short course of administration and MAM, a neurotoxin, is used for generating a neurodevelopmental model by administering the compound at the embryonic stage.

miR-146a-5p is significantly upregulated in MK-801 model whereas expression of miR-146a-3p exhibited a lesser extent of change and was not statistically significant (Figure 5A) (t=4.15, df=6 for miR-146a-5p; t=2.22, df=6 for miR-146a-3p). miR-200b-3p was significantly upregulated several folds in MK-801 model but the change in miR-200b-5p was not significant (Figure 5A) (t=6.6,df=7 for miR-200b-3p; t=1.42, df=7 for miR-200b-5p).

**Fig 5.**
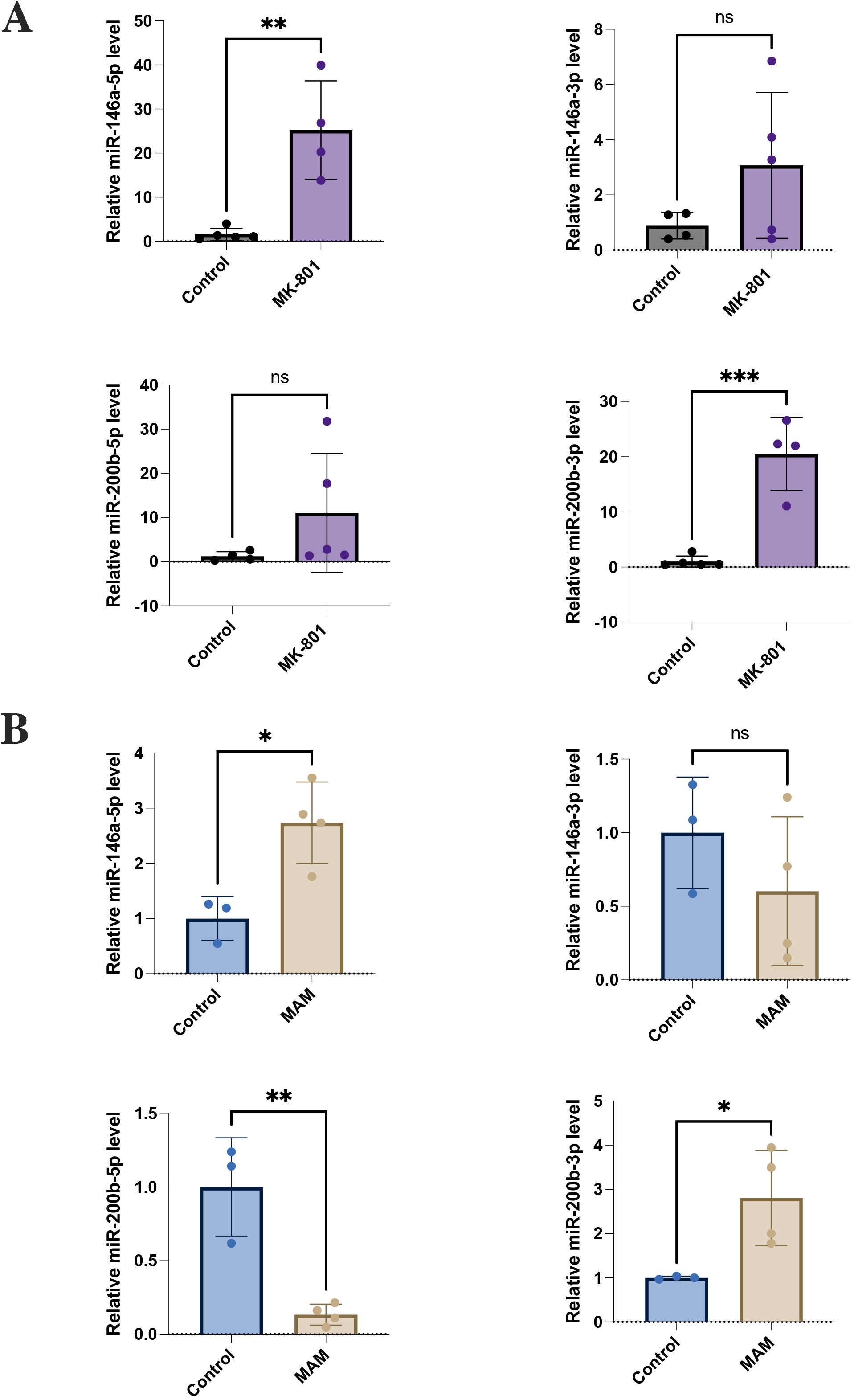
Upregulation of miR146a-5p and miR-200b-3p in MK-801 injectedand MAM injected animals. The MK-801 and MAM animal models were created as described before(Gunasekaran et al., 2021) and is mentioned in brief in the methods section (2.2.4.2.1).Real time PCR analysis was done for quantitating the levels of miR-146a-5p, miR-146a-3p, miR-200b-5p and miR-200b-3p in the MK-801 model (A) and the MAM model (B) in the hippocampal region. Data are expressed as mean ±SD. Statistical analysis was done using Student’s t test, ***p ≤ 0.001, **p ≤ 0.01, *p ≤ 0.05, n=3-5.

With MAM model, we observed significant upregulation of miR-146a-5p and miR-200b-3p (Figure 5B) (t=3.63, df=5 for miR-146a-5p; t=2.89, df=5 for miR-200b-3p). Interestingly we found significant downregulation in one of the complimentary forms, miR-200b-5p, but no significant change was observed in miR-146a-3p expression (Figure 5B) (t=5.19, df=5 for miR-200b-5p; t=1.21, df=5 for miR-146a-3p). These results showed that miR-146a-5p and miR-200b-3p were upregulated in both the models in which NMDAR subunits GluN2A and GluN2B were downregulated (Gunasekaran et al., 2021) consistent with a role of these miRNAs in causing decrease in the protein expression of GluN2A and GluN2B in these models.

## 4. DISCUSSION

Numerous studies have reported differential expression of miRNAs in the brain under various conditions but very few have addressed their physiological role in brain development and cognitive functions. Elucidating the specific roles of brain-enriched miRNAs would help in understanding neural mechanisms underlying brain functions and their importance in neurological disorders. This study examines the roles of miR-146a and miR-200b in regulating NMDAR subunits. These miRNAs were identified as candidates that can target *Grin2A* and *Grin2B*, by bioinformatics-assisted screening. Another major reason for choosing these miRNAs was their altered expression levels in neurological disorders (Wang et al., 2016; Mai et al., 2019; Liu et al., 2014;Fu et al., 2019; Beveridge et al., 2010; Perkins et al., 2007;Hall et al., 2015; Ghazaryan et al., 2019; Ibrahim et al., 2020; Moreau et al., 2011;Santarelli et al., 2011;). Using both *in vitro* and *in vivo* models, we showed interaction of the miRNAs with *Grin2A* and *Grin2B* and attenuation of their translation.We also found that in the two animal modelswhere GluN2A and GluN2B proteins were downregulated in the hippocampus, miR-146a-5p and miR-200b-3p were upregulated consistent with their involvement in downregulating NMDAR subunits.Complementary strands of miR-146a-3p and miR-200b-5p which do not target *Grin2A* and *Grin2B*, did not show a major change in expression thereby further supporting the mechanism (Figure 4A). We show severe learning and memory impairment upon overexpression of both the miRNAs *in vivo* (Figure 3) along with decreased levels of GluN2A and GluN2B protein expression (Figure 4C) indicating possible synaptic dysfunction.

Although, overexpression of miR-146a and miR-200b *in vivo* caused cognitive impairment, it did not lead to anxiety and depression related phenotype (Fig. 3C, Suppl. Fig. 3) indicating that upregulation of these miRNAs may only contribute to the cognitive component in these diseases.Our study suggests that the association of miR-146a and miR-200b to neurological disorderscould be, at least partly, due to their ability to downregulate NMDARs (Ghazaryanet al., 2019; Ibrahim et al., 2020; Fu et al., 2019) thusofferinga mechanistic explanation for diseases such as AD and schizophreniathathave an NMDAR component in their pathophysiology (Liu et al., 2019).

miR-146a and miR-200b, are known to affect and regulate other pathways. miR-146a is reported to also be associated with cerebrovascular diseases, neuroinflammation, CNS trauma, neuroautoimmune diseases, neuroviral infections, peripheral neuropathy, neurological tumor, ischemic stroke (IS), epilepsy and multiple sclerosis (Fan et al., 2020, Shomali et al., 2017; Juźwik et al., 2019). It has roles in neuronal survival and axonal growth (Zhou et al., 2016; Jia et al., 2016) and in development by targeting NFκB, NOTCH and WNT/β-catenin (Taganov et al., 2006; Huang et al., 2016; Ghahhari and Babashah, 2015; Hwang et al., 2012). Numerous studies showed miR-200 family to be associated with thepathogenesis of neurodegenerative diseases such as Parkinsons disease (PD), ALS and Huntingtons disease (HD) (Fu et al., 2019). A recent report suggests a role for miR-200b-3p in hypoxia-ischemia brain damage by upregulating Slit2 expression. Inhibiting miR-200b-3p alleviates brain injury in neonatal rat model of hypoxia-ischemic brain damage (HIBD) (Zhang et al., 2020). miR-200 family is found toplay key roles in the control of epithelial-mesenchymal transition (EMT) in neurogenesis (Beclin et al., 2016) and in processes such as initiation, progression and metastasisin gliomas (Peng et al., 2018).

Interestingly, we obtained data on inverse correlation of miR-200b-5p and miR-200b-3p levels in the MAM treatment model (Figure 6). This could be due to two mechanisms which are recently gaining attention-target mediated miRNA protection (TMMP) and/or target RNA directed miRNA degradation (TDMD) (Ameres et al., 2010; Haas et al., 2016; de la Mata et al., 2015).

**Fig 6.**
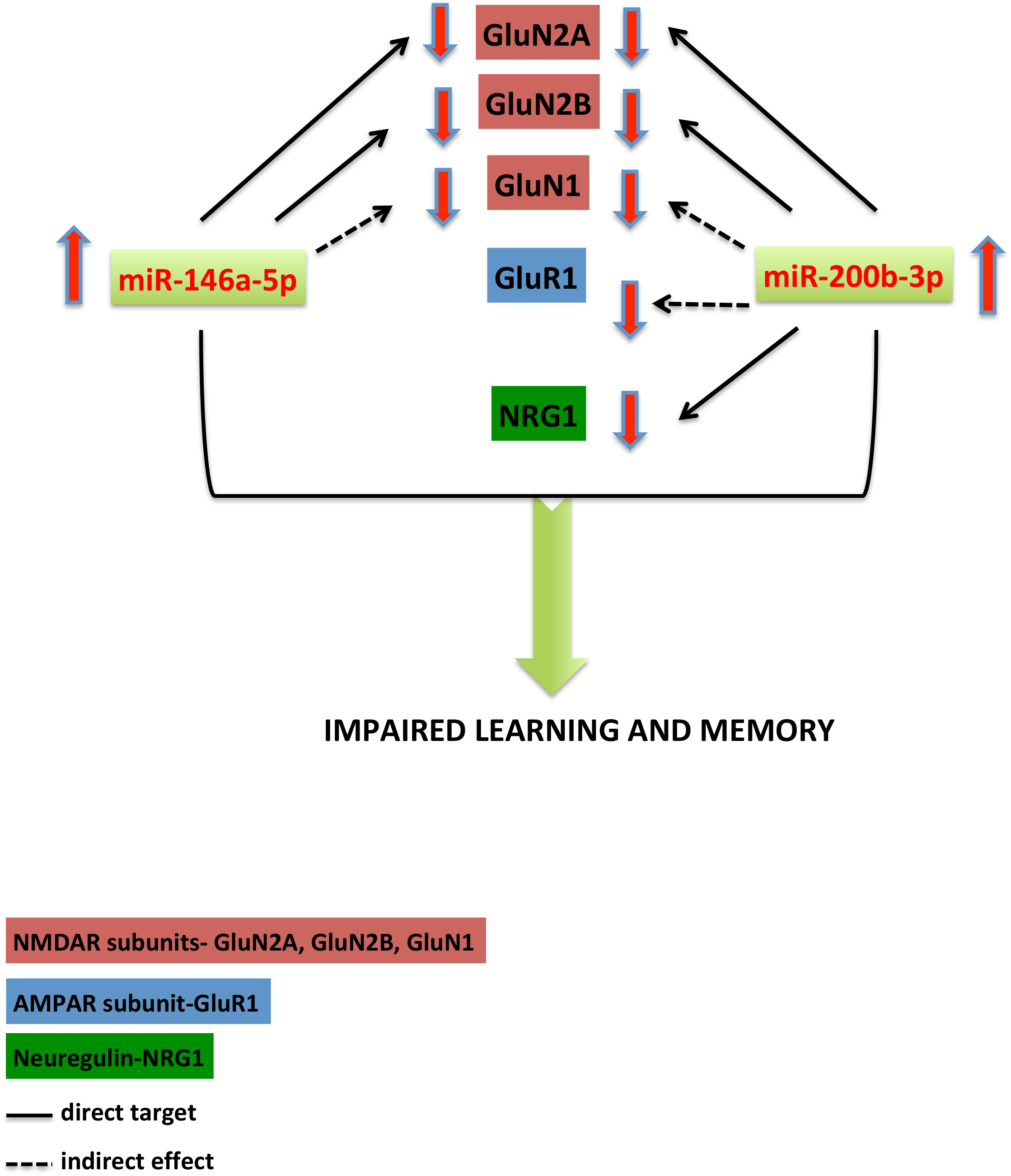
Schematic representation of the downregulation of proteins by the overexpressed miR-146a-5p and miR-200b-3p. Overexpression of miR-146a-5p or miR-200b-3p in rats leads to downregulation of GluN1, GluN2A and GluN2B subunits of NMDAR with GluR1 andNRG1 only in miR-200b-3p injected animals thus affecting cognition

MiRNAs are posttranscriptional regulators which bind to their target messenger RNAs (mRNAs) resulting in mRNA destabilisation and translational repression (O’Brien et al., 2018). But there are condratictory reports on decreased mRNA levels with elevated miRNA expression. Hill et al proposed a regulatory network model in which miRNAs might not profoundly change the target mRNA levels. It states that the downregulation of the target mRNA might be masked because of increased mRNA transcription or by downregulation of its repressor genes that are targeted by the same miRNAs (Hill et al., 2014). Our data shows that the mRNA levels were not affected in the animals in which miRNAs were overexpressed in the hippocampal region (Figure 5B). The protein levels were significantly reduced for these transcripts indicating that translational repression without mRNA cleavage would be the mechansim employed in downregulating NMDARs under these conditions (Figure 4B and 4C).

We also checked for GluN1 protein levels as it is one of the major subunits of NMDAR. There was reduction in GluN1 protein which might be a secondary effect of reduced GluN2A/GluN2B protein levels(Figure 2D and 4C)as GluN1 is not a direct target of these miRNAs based on bioinformatic analysis. Because of the persistent decrease in the levels of GluN2A/GluN2B proteins, the functional NMDAR heteromer, GluN1/GluN2, would be low leading to impairment in learning and memory tasks. The majority of native NMDA receptors are tetrameric assemblies of two GluN1 subunits and two GluN2 subunits (Traynelis et al., 2010), in particular GluN1/GluN2A/GluN2B triheteromers. This combination accounts for 50% of the total NMDAR in hippocampus and cortex of an adult rodent brain (Hansen et al., 2014). GluN1/GluN2/GluN3 heteromers are also present but their relative abundance is unknown. From our data, we can extrapolate that miR-146a and miR-200b can independently downregulate GluN1/GluN2A/GluN2B triheteromers which forms a major proportion of NMDARs leading to significant functional alterations *in vivo*. It will be relevant to investigate the role of these miRNAs in synaptic plasticity mechansims that are likely to be hampered in our miRNA overexpression models.

NRG1 was downregulated by miR-200b-3p as predicted by bioinformatic analysis (Suppl. Figure 4). This was observed in primary hippocampal neurons and in AAV-miR-200b injected animals (Figure 2J and 4C). There are two sites of interaction in*NRG1* 3’UTR and both are highly conserved among vertebrates (Suppl. Fig 4)with site type 7mer-m8. miR-146a-5p was also predicted to show interaction with *NRG1* at sites conserved only in rat and mouse, but there was no significant effect on the protein level in our experiments *in vitro* and *in vivo* in which miR-146a was overexpressed. We also observed decrease in the GluR1 levels in miRNA injected animals (data statistically significant for only AAV-miR-200b injected animals) (Figure 2I and 5C). GluR1 is an important AMPA receptor subunit playing crucial roles in synaptic plasticity (Chater et al., 2014). Decrease in GluR1 could be a secondary effect of either NMDAR downregualtion or overexpression of these miRNAs, as there are no direct binding sites for miR-200b to *Gria1*.

Understanding the role of miRNAs will provide insights on how to manipulate the levels of certain proteins which are altered in neurological disesaes. Manipulating NMDAR activity as such leads to severe side effects in patients (Lipton et al., 2004). Hence alterative theraputic stratergies such as using miRNAs could play a significant role in modulating these proteins without much side effects. Our study provides insights on how upregulated miR-146a-5p and miR-200b-3p can contribute tosevere cognitive impairment inneurological disorders (Figure 6).

## 5. CONCLUSION

Our objective was to identify miRNAs that target the NMDA receptors and study their role in learning and memory. For this we selected miR-146a and miR-200b, which were differentially expressed in neuropsychiatric and neurodegenerative disorders. By bioinformatics analysis it was found that the two miRNAs have target sites on the transcripts of the NMDAR subunits GluN2A and GluN2B. Interaction of these miRNAs with *Grin2A* and *Grin2B* was shown by luciferase assay. Western blot analysis of primary hippocampal neurons overexpressed with these miRNAs showed downregulation of GluN2A and GluN2B. Overexpression of these miRNAs *in vivo* in the hippocampus of rats through stereotactic injection of AAV particles caused downregulation of GluN2A, GluN2B and GluN1 proteins without significant alterations in their transcript levels. Additionally, protein levels of NRG1 and GluR1 were also significantly downregulated in AAV-miR-200b injected animals. Both the miRNAs caused learning and memory defects upon overexpression in animals as expected from the molecular level changes. The levels of miR-146a-5p and miR-200b-3p were also upregulated in two rat models in which the NMDAR subunits were downregulated. These results highlight the key roles of small non-coding RNAs in modulating NMDAR and learning and memory processes thus bringing out the possibility of using them in therapeutic interventions.

## Abbreviations

NMDAR: N-methyl-D-aspartate receptor
MAM: Methylazoxymethanol acetate
miRNA: microRNA
Pre-miRNA: precursor miRNA
AAV: Adeno-associated virus
VC: Vector control
AAV-miR-200b: GFP-rno-mir-200b AAV miRNA virus
AAV-miR-146a: GFP-rno-mir-146a AAV miRNA virus
NRG1: Neuregulin1
3’UTR: 3’ Untranslated region
CaMKII: Calcium/calmodulin dependent protein kinase II
GluN2B/*Grin2B*: 2B subunit of NMDAR
GluN2A/*Grin2A*: 2A subunit of NMDAR
NORT: Novel object recognition test
OLT: Object location test
OFT: Open field test
MWM: Morris water maze
RI: Recognition index
DI: Discrimination index

## Acknowledgements

We are grateful to Dr. Ani V Das and Dr. Jackson James for providingluciferase plasmid psiCHECK2 (Promega) and pRIPM plasmid and for assistance. We thank Ms. Reena Sarah Jacob for performing the statistical analysis. We thank Mr. K.C. Sivakumar for technical advice on bioinformatics. We thank Dr. Mayadevi, Dr. Mantosh Kumar and Dr. Lakshmi K for their support and advice. We also thank the funding agencies-RGCB and DST, India.

## Statements & Declarations

### Funding

This work has been supported by Rajiv Gandhi Centre for Biotechnology and Department of Science and Technology (DST), Government of India.

### Author Contributions

SG and RVO contributed to conception and design of the study. SG and RVO developed the methodology. SG performed the experiments. RVO acquired and provided resources and overall supervision for the study. SG wrote the first draft of the manuscript. Both the authors contributed to manuscript revision and approved the submitted version.

### Conflicts of Interest

The authors declare no competing interests.

### Ethical Statement

All procedures for animal experiments followed the rules and regulations prescribed by the Committee for the Purpose of Control and Supervision of Experiments on Animals (CPCSEA), Government of India and were approved by the Institutional Animal Ethics Committee (IAEC) of Rajiv Gandhi Centre for Biotechnology.

### Consent to Participate

Not applicable

### Consent for Publication

All authors have approved the contents of this manuscript and provided consent for publication.

### Data Availability Statement

Data supporting the results is provided in the manuscript and also is available with the authors.

## Notes

### Competing Interest Statement

The authors have declared no competing interest.

