## Supplementary Information for "miR-146a and miR-200b alter cognition by targeting NMDA receptor subunits"

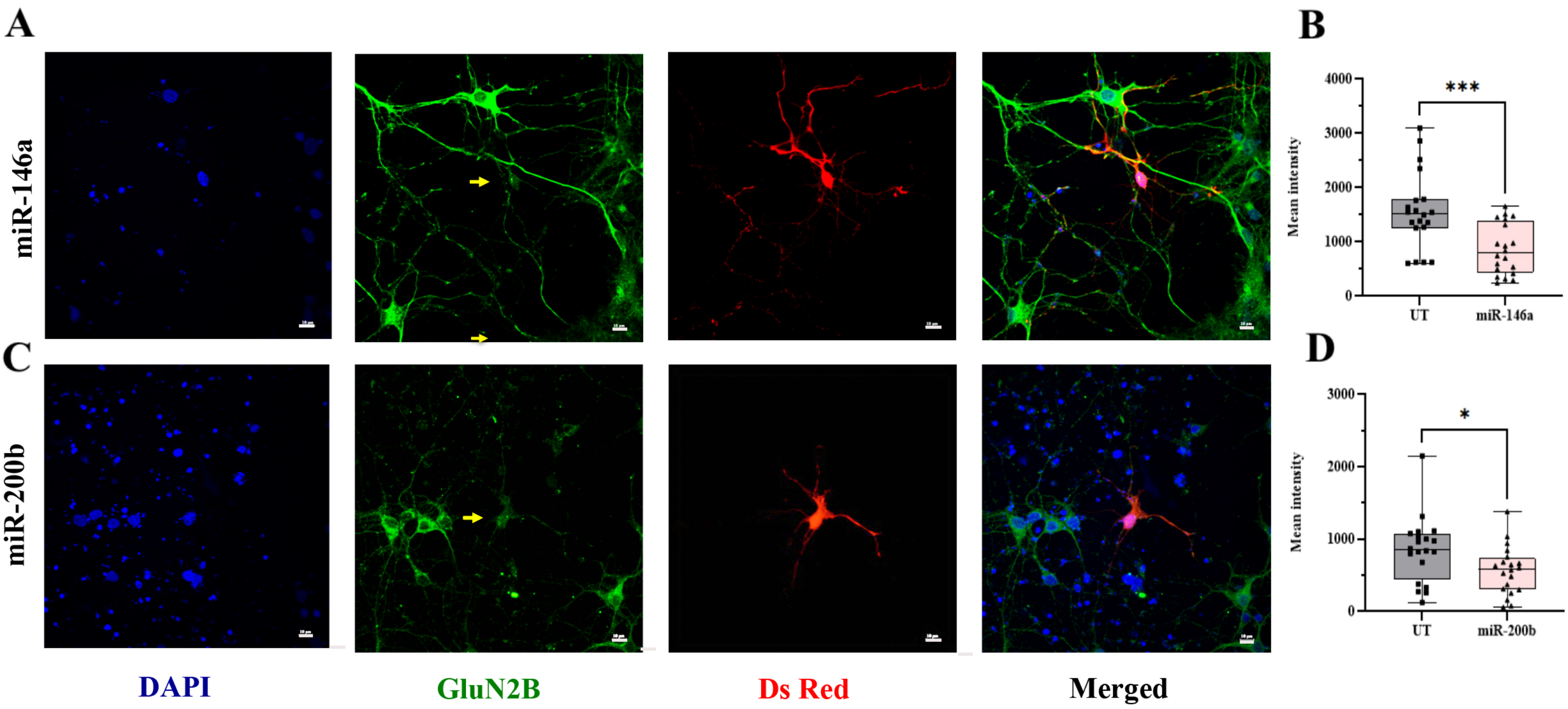


**Suppl. Fig 1 Downregulation of GluN2B protein upon overexpression of miR-146a and miR-200b.**
(A and C) Pre-miRNA-146a and pre-miR-200b sequences present in pRIPM vector were transfected in primary hippocampal neurons on DIV 7 and the cells were fixed on DIV 10 using 4% paraformaldehyde. The cells were washed, blocked and were incubated with primary anti-GluN2B antibody (rabbit, 1:500, Abcam) overnightfollowed by secondary antibody incubation for 2 hrs using fluorophore conjugated secondary antibody (Alexa Fluor 488, green) along with DAPI stainingfor nucleus (blue). Images were captured using Nikon confocal microscope (Nikon EclipseTi A1R). Images are representative of 3 independent experiments(B and D). Using NIS Elements V.4.0 imaging software, the region of interest (ROI) quantification was performed on untransfected (UT) and miRNA transfected cells for GluN2B expression. Most of the fields had around 10-12 untransfected cells with 1-2 transfected cells. For each field [(Total no. of fields quantified in 3-4 experiments)=20], the ROI quantification values were averaged for untransfected and transfected cells. Data presented are mean ± SD. Boxes indicate the lower quartile, median and upper quartileand whiskers indicate the range of changes (minimum to maximum). Statistical analysis was done by Student's t test, ***$p \leq 0.001$, *$p \leq 0.05$, n=3

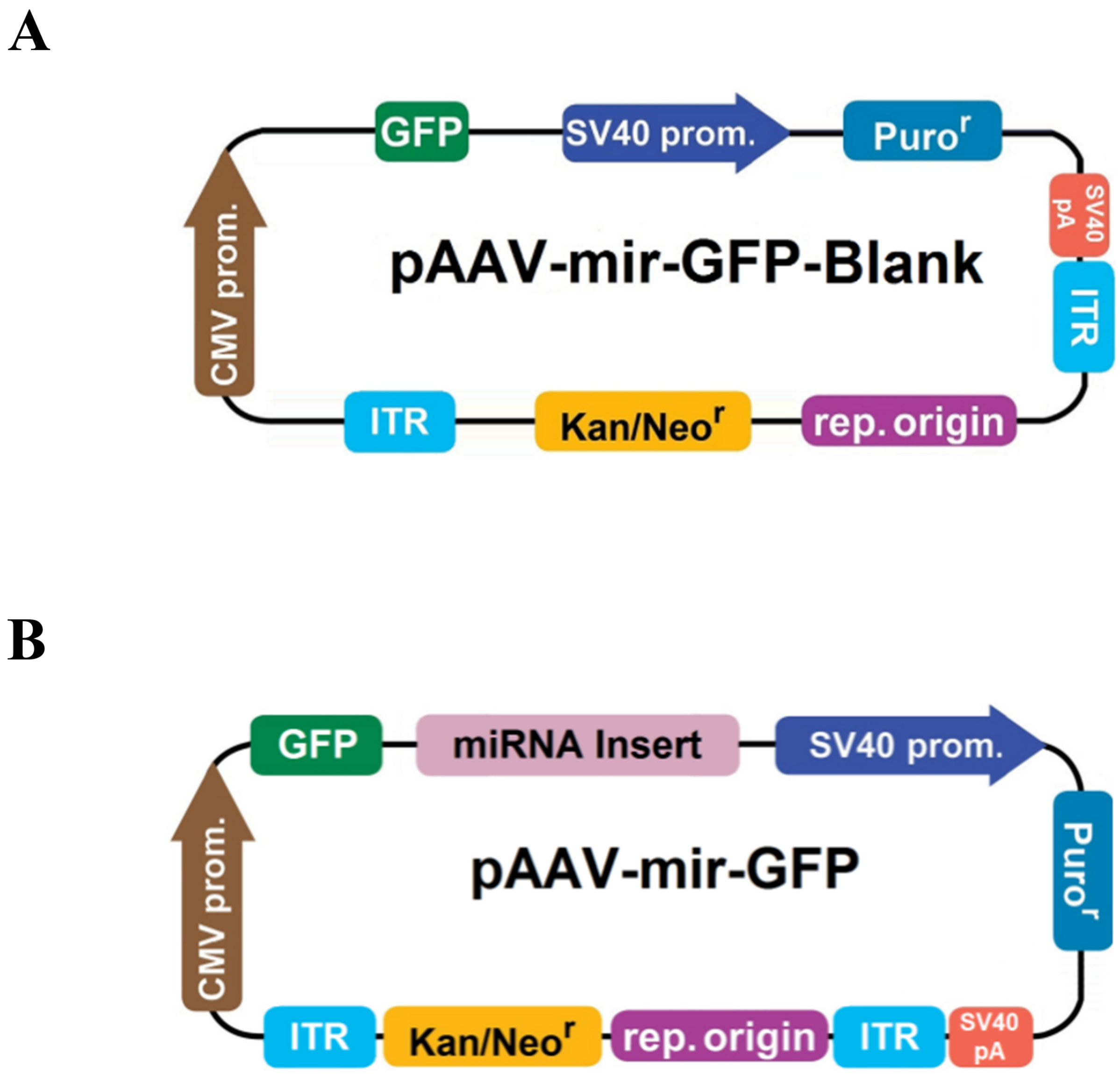


**Suppl. Fig 2 The vector backbone used for the production of AAV particles.** The plasmid maps of (A) vector control and (B) miR-146a and miR-200b

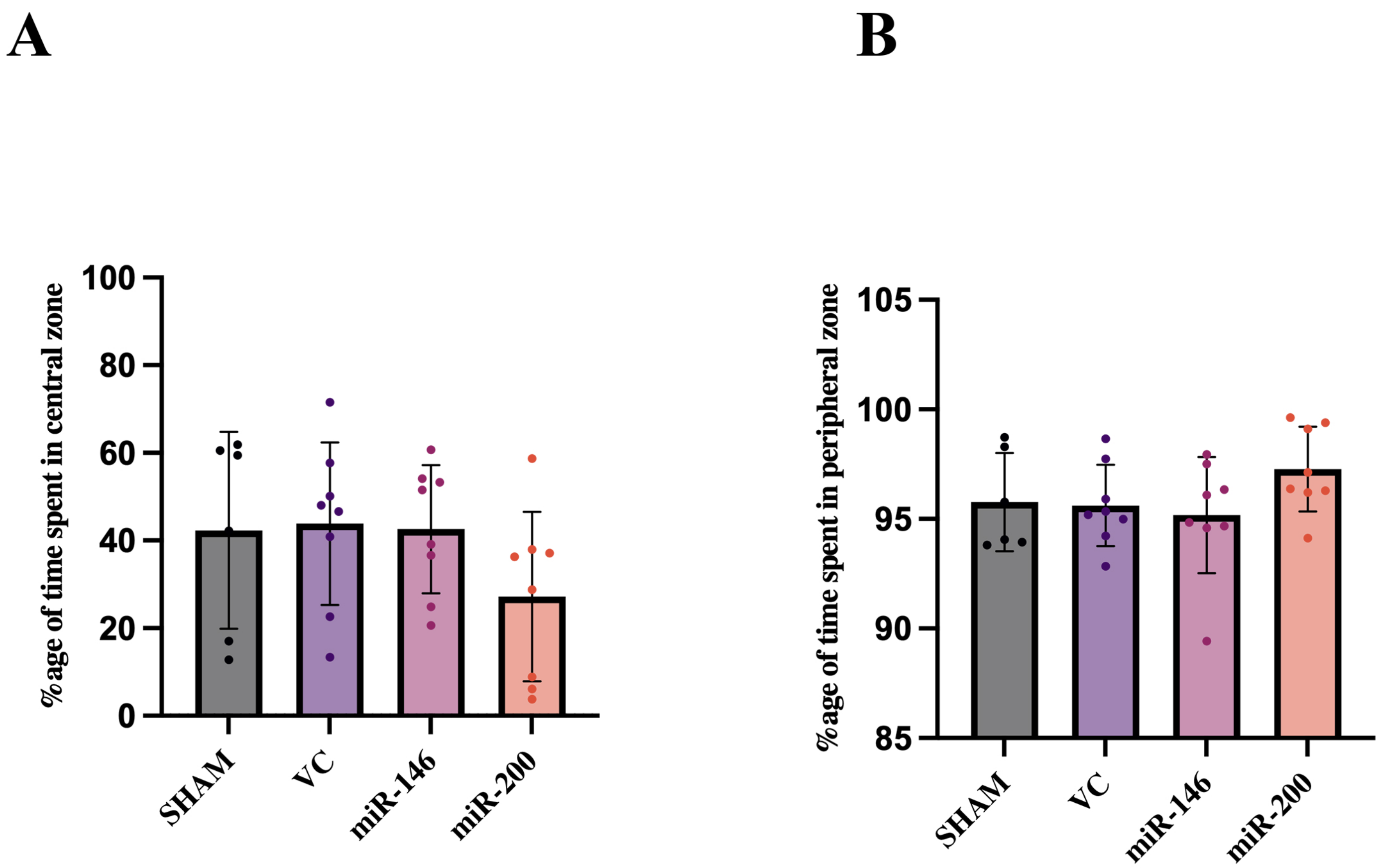


**Suppl. Fig 3 The animals injected with miR-146a and miR-200b show no significant changes in anxiety levels.** Open field test showing the time spent in the (A) central zone and (B) peripheral zone during the total period of 10-11 min. Data is shown as mean± SD; n=6-8 per group

**A**

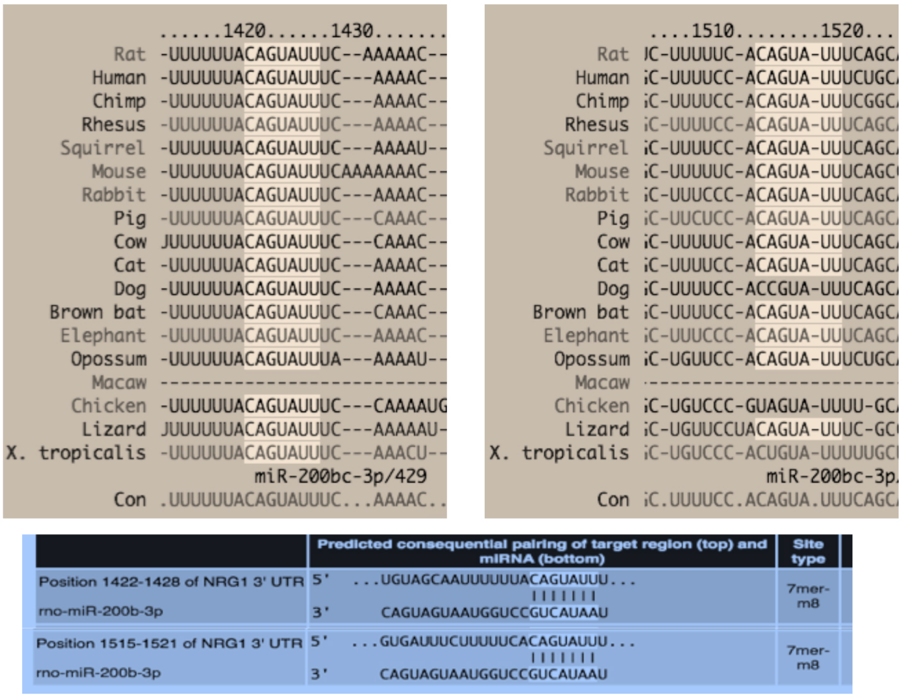


**B**

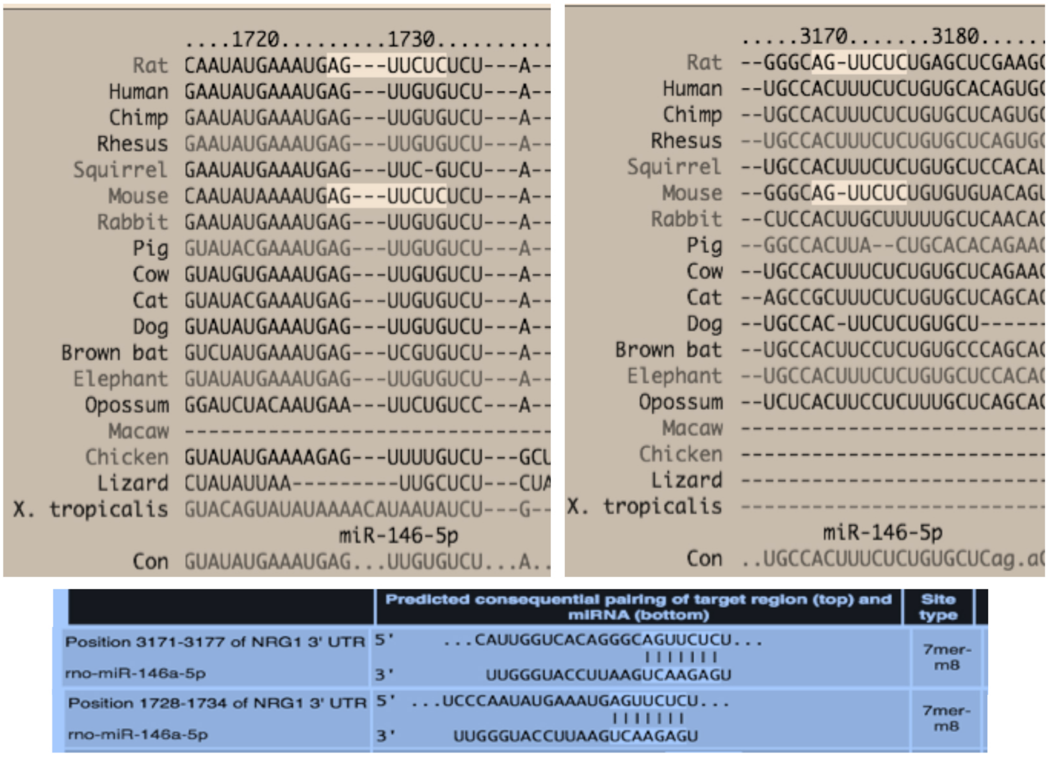


**Suppl. Fig 4 Sequence alignment of miR-146a and miR-200b with Neuregulin 1 (NRG1).** (A) Bioinformatic analysis results showing conserved miR-200b target sequence within NRG1 at two positions (B) Conservation of miR-146a target sequence within NRG1 between rat and mouse.
